# How thermophilic Gram-positive organisms perform extracellular electron transfer: characterization of the cell surface terminal reductase OcwA

**DOI:** 10.1101/641308

**Authors:** N.L. Costa, B. Hermann, V. Fourmond, M. Faustino, M. Teixeira, O. Einsle, C.M. Paquete, R.O. Louro

**Affiliations:** Instituto de Tecnologia Química e Biológica António Xavier, Universidade NOVA de Lisboa, Portugal; Institut für Biochemie, Albert-Ludwigs-Universität Freiburg, Germany; Aix-Marseille Université, Centre National de la Recherche Scientifique (CNRS), France; BIOSS Centre for Biological Signaling Studies, Albert-Ludwigs-Universität Freiburg, Germany

**Author notes:** Corresponding authors: Oliver Einsle, Catarina M. Paquete. Contributed equally to the contents of the manuscript (N.L. Costa, B. Hermann).

## Abstract

Extracellular electron transfer is the key process underpinning the development of bioelectrochemical systems for the production of energy or added-value compounds. *Thermincola potens* JR is a promising Gram-positive bacterium to be used in these systems because it is thermophilic. In this paper we describe the structural and functional properties of the nonaheme cytochrome OcwA, which is the terminal reductase of this organism. The structure of OcwA, determined at 2.2Å resolution shows that the overall-fold and organization of the hemes are not related to other metal reductases and instead are similar to that of multiheme cytochromes involved in the biogeochemical cycles of nitrogen and sulfur. We show that, in addition to solid electron acceptors, OcwA can also reduce soluble electron shuttles and oxyanions. These data reveal that OcwA can take the role of a respiratory ‘swiss-army knife’ allowing this organism to grow in environments with rapidly changing availability of terminal electron acceptors without the need for transcriptional regulation and protein synthesis.

**Importance:** Thermophilic Gram-positive organisms were recently shown to be a promising class of organisms to be used in bioelectrochemical systems for the production of electrical energy. These organisms present a thick peptidoglycan layer that was thought to preclude them to perform extracellular electron transfer (i.e. exchange catabolic electrons with solid electron acceptors outside of the cell). In this manuscript we describe the structure and functional mechanisms of the multiheme cytochrome OcwA, the terminal reductase of the Gram-positive bacterium *Thermincola potens* JR found at the cell surface of this organism. The results presented here show that this protein is unrelated with terminal reductases found at the cell surface of other electroactive organisms. Instead, OcwA is similar to terminal reductases of soluble electron acceptors. Our data reveals that terminal oxidoreductases of soluble and insoluble substrates are evolutionarily related, providing novel insights into the evolutionary pathway of multiheme cytochromes.

## Introduction

Iron is one of the most abundant metals in Earth’s crust, and microbial reduction of iron is associated with some of the earliest life forms (1). Nowadays, this type of metabolism is used for numerous biotechnological processes, including production of energy in microbial fuel cells (MFC) and synthesis of added-value compounds in microbial electrosynthesis (MES) (2, 3). It is the ability to perform extracellular electron transfer that allows some microorganisms to exchange electrons with an electrode in these devices (4). While microbes in MFCs donate electrons to electrodes and generate electrical current (3), in MES the electrode oxidation performed by bacteria is coupled to the production of chemicals on the cathode compartment (5). Currently, about 100 microorganisms are known to perform extracellular electron transfer and exchange electrons with an electrode (6). Most of these organisms are Gram-negative bacteria (6); this is mainly because of the long-held view that the thick peptidoglycan layer that encases Gram-positive bacteria prevents them from performing this type of metabolism (7). However, recently it was described that Gram-positive bacteria are also able to perform extracellular electron transfer (8, 9). The iron-reducing Gram-positive bacterium *Thermincola (T.) potens* JR was identified in a current-producing MFC operating at high temperature (10), whereas the closely related bacterium *T. ferriacetica* was isolated from ferric deposits in a hydrothermal spring (11). These two thermophilic microorganisms (10) were shown to produce higher current levels in MFC when compared with mesophilic organisms in the same type of bioreactor (10, 12). Despite their faster kinetics and lower interference from oxygen intrusion (12, 13), their application is still hindered by the lack of knowledge of the molecular mechanism that they use for extracellular electron transfer. This has a negative impact in the ability to optimize bioelectrochemical systems for the efficient production of energy using this promising class of organisms.

Like Gram-negative electroactive organisms, *Thermincola* sp. contain a large number of genes that code for multiheme *c*-type cytochromes (MHCs) (10, 14). Recently, a putative electron transfer pathway was proposed for *T. potens* JR (8). In this pathway, the nonaheme cytochrome TherJR_2595 (Tfer_3193 in *T. ferriacetica*) is located at the cell surface and was proposed to be the terminal reductase for extracellular electron transfer (8). The characterization of this protein is essential to elucidate the molecular mechanisms of electron transfer at the electrode-microbe interface in Gram-positive bacteria.

In this work, we describe the structural and functional properties of TherJR_2595, hereafter referred to as OcwA (<u>O</u>uter <u>c</u>ell-<u>w</u>all protein <u>A</u>). The three-dimensional structure of this protein reveals that it is unrelated to the structurally characterized outer-membrane MHCs from *Shewanella*. Instead, it is structurally and functionally related with MHCs involved in the biogeochemical cycles of nitrogen and sulfur, which suggests that terminal reductases that use soluble and insoluble electron acceptors may be evolutionarily related. This work provides the first insight into the molecular mechanisms of terminal reductases from thermophilic Gram-positive bacteria, which underpin the direct and indirect electron transfer to electrodes. This knowledge is crucial for the implementation of MFC with a broader microbiological range and at more varied operational conditions, such as at high temperatures.

## Results

### Production of OcwA

Recombinant OcwA was purified to electrophoretic homogeneity and identified as a band at approximately 62 kDa in SDS-PAGE (Figure 1A). This band stained positively for covalently attached hemes (Figure 1A), and N-terminal sequencing retrieved the predicted sequence of OcwA (EKPAD) without the signal peptide, showing that the protein was efficiently processed in *E. coli*. Size exclusion chromatography of pure OcwA revealed an approximately equal mixture of monomer and dimer forms (Figure SM1). UV-visible spectra of OcwA showed the typical features of a low-spin cytochrome *c* (Figure 1B), and NMR and EPR spectroscopy indicated that this protein contains at least three types of hemes (Figure 2).

**Figure 1.**
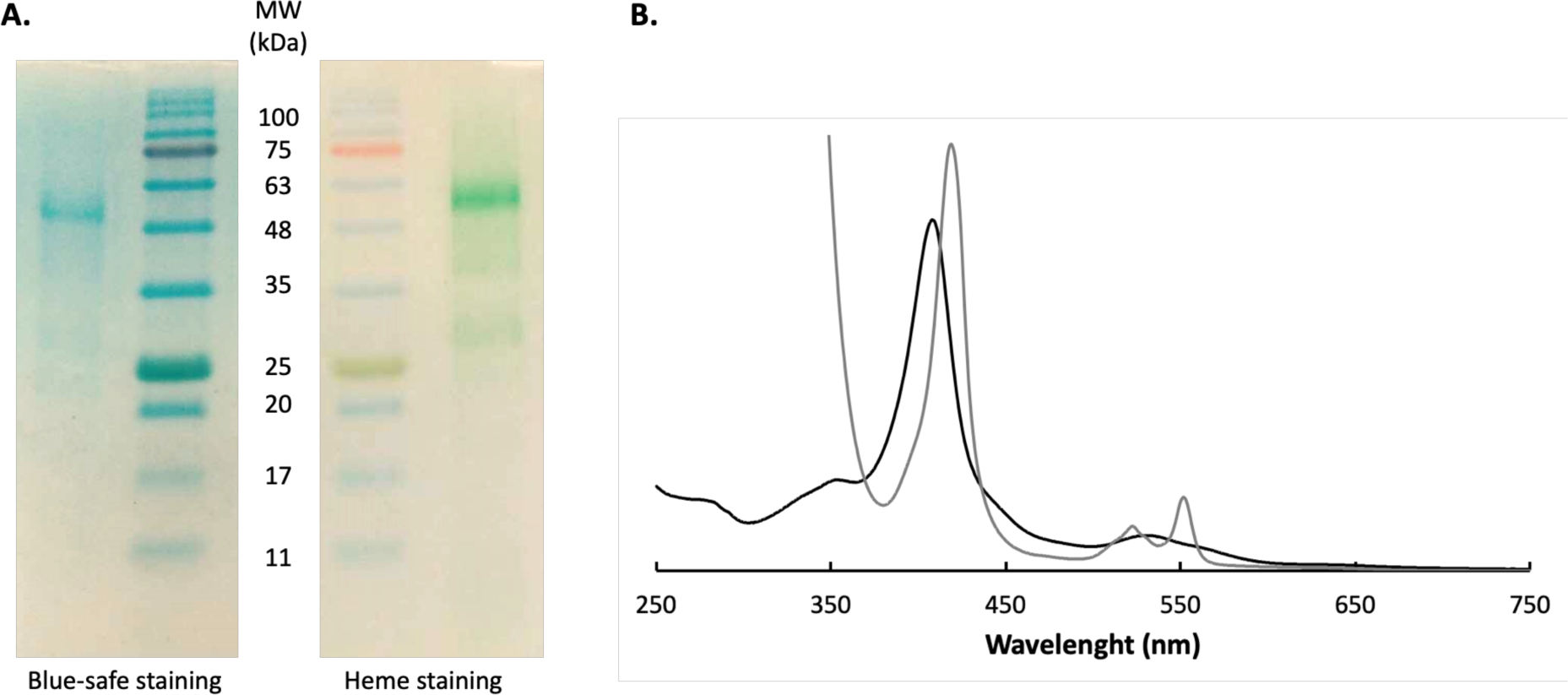
Isolation of OcwA from *T. potens* JR. **A.** Blue-safe and heme stained SDS-PAGE of purified OcwA. **B.** UV-visible spectra of OcwA obtained in the reduced (grey) and oxidized state (black).

**Figure 2.**
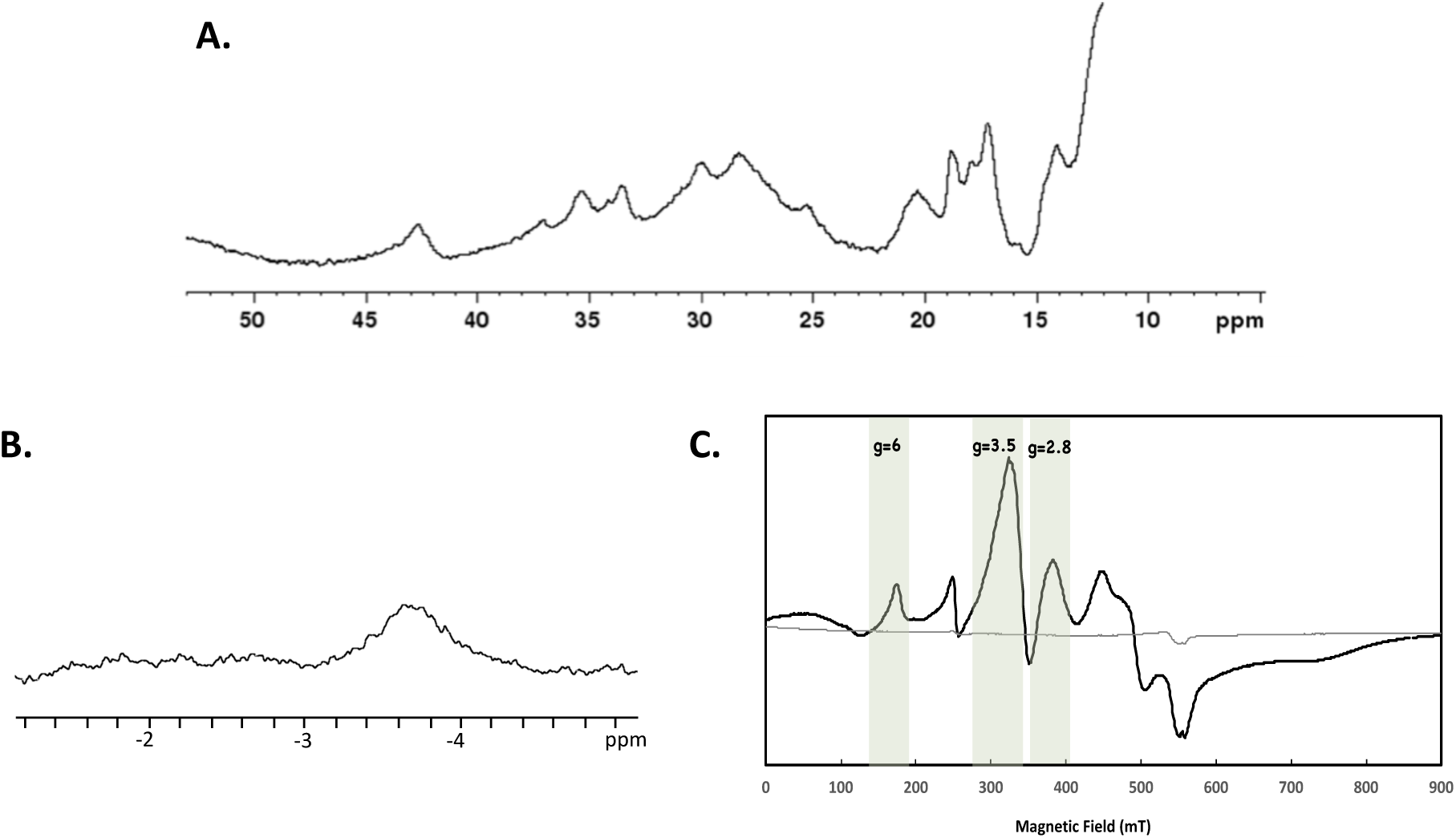
Magnetic spectroscopic properties of OcwA from *T. potens* JR. **A.** ^1^H NMR spectrum of OcwA obtained at 25°C in the oxidized state **B.** 1D ^1^H NMR spectrum of OcwA in the reduced state obtained at 25°C by the addition of sodium dithionite. **C.** EPR spectra of OcwA at 9.39GHz in the oxidized (black line) and reduced (grey line) state at 7K. The unlabeled signal at g=4.3 likely corresponds to a small amount of high spin ferric iron adventitiously present in the sample.

The ^1^H NMR spectrum of OcwA in the oxidized state presents signals outside of the protein envelope up to 45 ppm (Figure 2A). These signals display a Curie-type temperature dependence (Figure SM2), which are typical of the methyl groups of low-spin ferric (Fe^3+^) hemes with two strong-field axial ligands such as histidines or methionines (15). Reduction of OcwA with sodium dithionite revealed an NMR peak near −3 ppm (Figure 2B). This indicates that besides the typical bis-histidine axial coordinated hemes, OcwA also contains at least one heme that is axially coordinated by a methionine (16). Interestingly, this occurs even though the amino acid sequence of OcwA contains enough histidines for all the nine hemes to be bis-histidine coordinated.

Continuous-wave X-band EPR spectroscopy shows that reduced OcwA is silent whereas the oxidized protein revealed a superposition of signals that suggest the presence of at least three groups of paramagnetic species (Figure 2C). The shape of the spectrum shows clearly the presence of magnetic interactions among the hemes, as often observed in MHCs and expected due to the short distance between hemes in OcwA (as detailed in next section). The signal with *g*_max_=2.8 is characteristic of low-spin (S=1/2) six-coordinated hemes with axial ligands approximately parallel with each other, whereas signals with *g*_max_=3.5 are due to low-spin six-coordinated hemes with axial ligands perpendicular to each other (17). The resonance at *g* ∼ 6 indicates the presence of high-spin hemes (S=5/2) that are typically five-coordinated.

### 3D-structure of OcwA

The three-dimensional structure of OcwA was determined to 2.2 Ả resolution by X-ray crystallography (Table SM1). OcwA is composed by a globular heme domain and a three-helix bundle at the C-terminal. These three long α-helices are reminiscent of the pentaheme NrfA (nitrite reductase) family of proteins (18) and of the octaheme cytochromes hydroxylamine oxidoreductase (HAO) and (19) sulfite reductase MccA (Figure 3A) (20). The crystal structure of OcwA shows that this protein forms a dimer, as observed experimentally by gel filtration (Figure SM1). It spans about 180 × 81 × 45 Å, but presents only limited contact at the interface between monomers involving two regions. While the first contact area is formed by the base – but not the remainder – of the three-helix bundle, the second is made by a small alpha helix and a neighboring loop from an additional beta-sheet domain (amino acids 75-156) within the globular heme domain. This is an unprecedented monomer arrangement, where the three-helix bundle does not dominate the formation of the dimer interface. This arrangement may be a consequence of the decreased length of the C-terminal helices with respect to NrfA proteins.

**Figure 3.**
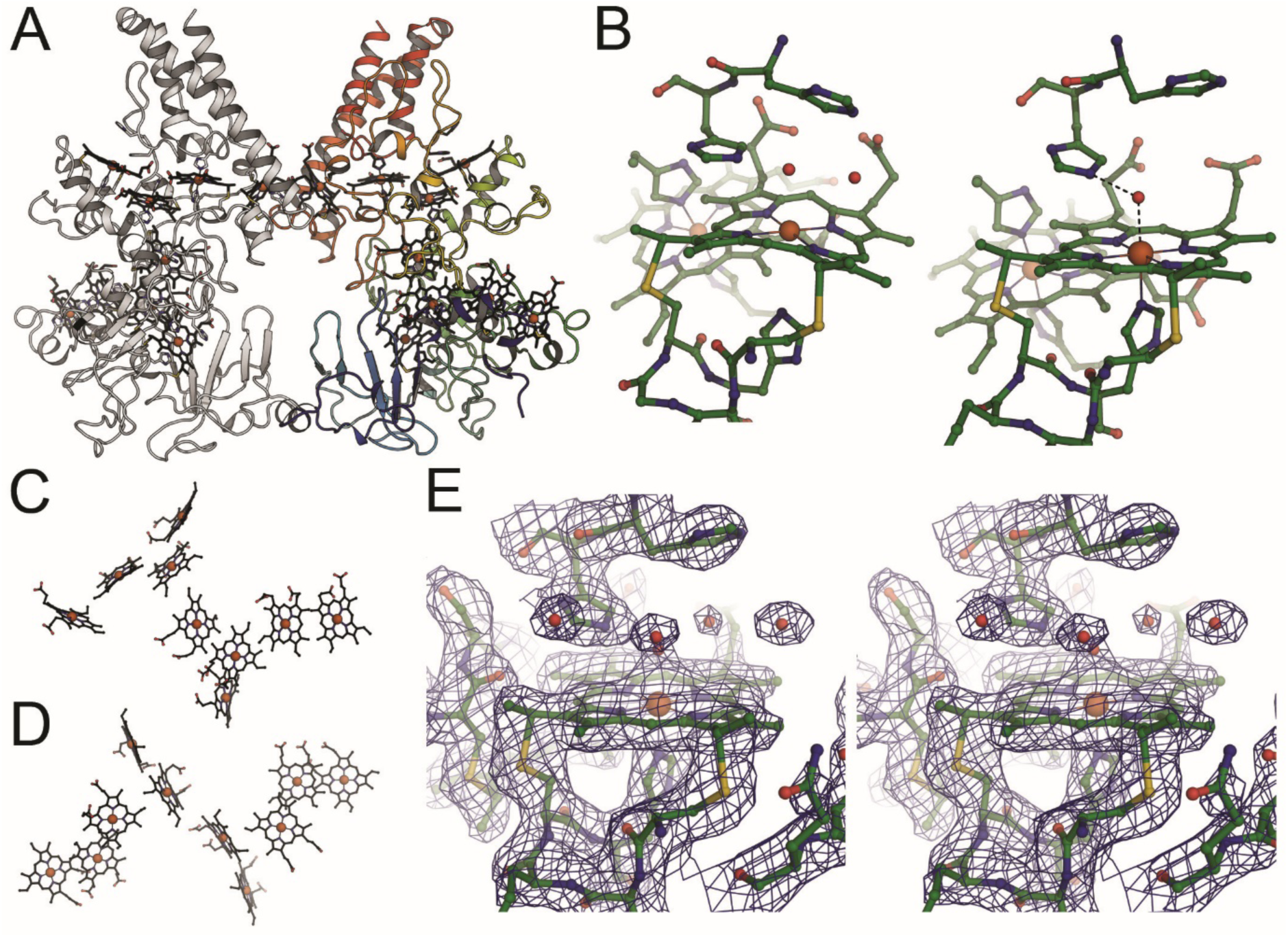
Structure of OcwA from *T. potens* JR. **A.** Dimer structure of OcwA. The right monomer is colored from blue at the N-terminus to red at the C-terminus. Heme groups are depicted in stick representation. **B.** Close-up of the active site heme groups 2 (left) and 5 (right) highlighting the similar environment of both centres, and depicting the water molecules found on the distal side. Comparison of the heme cores of **C**. OcwA and **D.** MtrC (PDB 4LM8). **E.** Stereo representation of the 2*F*_o_–*F*_c_ electron density map surrounding heme group 2, contoured at the 1 σ level.

The hemes in OcwA present three distinct coordination environments: hemes 1, 3, 4, 6, 7 and 8 (numbered according to the position of the CXXCH binding motif in the polypeptide sequence, where X is any amino acid) are bis-histidine coordinated. Among these, hemes 1, 3 and 6 likely contribute to the EPR signal with g= 2.8 because their axial histidines have nearly parallel rings. By contrast, hemes 4, 7 and 8 likely contribute to the EPR signal with g= 3.5 as their axial histidines have nearly perpendicular rings. Heme 9 features a histidine-methionine coordination, whereas hemes 2 and 5 have histidine as a proximal ligand and an open coordination site at the distal position (Figure 3B), being thus responsible to the EPR signal with g=6. These two hemes, located at opposite ends of the nine-heme arrangement, have an almost identical environment, with a histidine serving as proximal iron ligand and a His-His motif (HH 281/282 at heme 2 and HH 380/381 at heme 5) situated above the distal face of the heme, which in both cases forms a solvent-exposed pocket. The presence of these two penta-coordinated hemes that work as two putative active sites for substrate binding within a single monomer is a novelty within the family of MHCs (21).

The organization of the hemes within OcwA, although vaguely reminiscent of the ‘staggered-cross’ design of the structurally characterized outer-membrane MHCs (22) (Figure 3C), clearly follows a different design, that is similar to that of the NrfA family of proteins, with alternating parallel and perpendicular di-heme packing motifs. In fact, hemes 1-4 of OcwA align with a root-mean-squared deviation (rmsd) for all atoms of 0.84 Å with the hemes of the tetraheme cytochrome *c*_554_ from *Nitrosomonas europaea.* In both proteins, heme 2 is the active site (Figure 4B). Moreover, hemes 5-9 of OcwA can be superimposed with a rmsd of 0.89 Å to the NrfA heme core structure (Figure 4C), with heme 5 of OcwA as the second active site. Interestingly, hemes 1-4 and 6-9 also align with a rmsd of 0.89 Å to the heme core structure of the sulfite reductase MccA (20), with heme 2 in both proteins acting as the active site (Figure 4D). Although there is some correspondence of the polypeptides of cytochrome *c*_554_, NrfA and MccA with the corresponding sections of OcwA, sequence similarity is only observed for the heme-binding motifs, the distal heme ligands except for those of heme 2 and the three-helix bundle. Furthermore, MccA and OcwA share the additional β-sheet domain that has no equivalence in other NrfA family proteins (Figure SM3).

**Figure 4.**
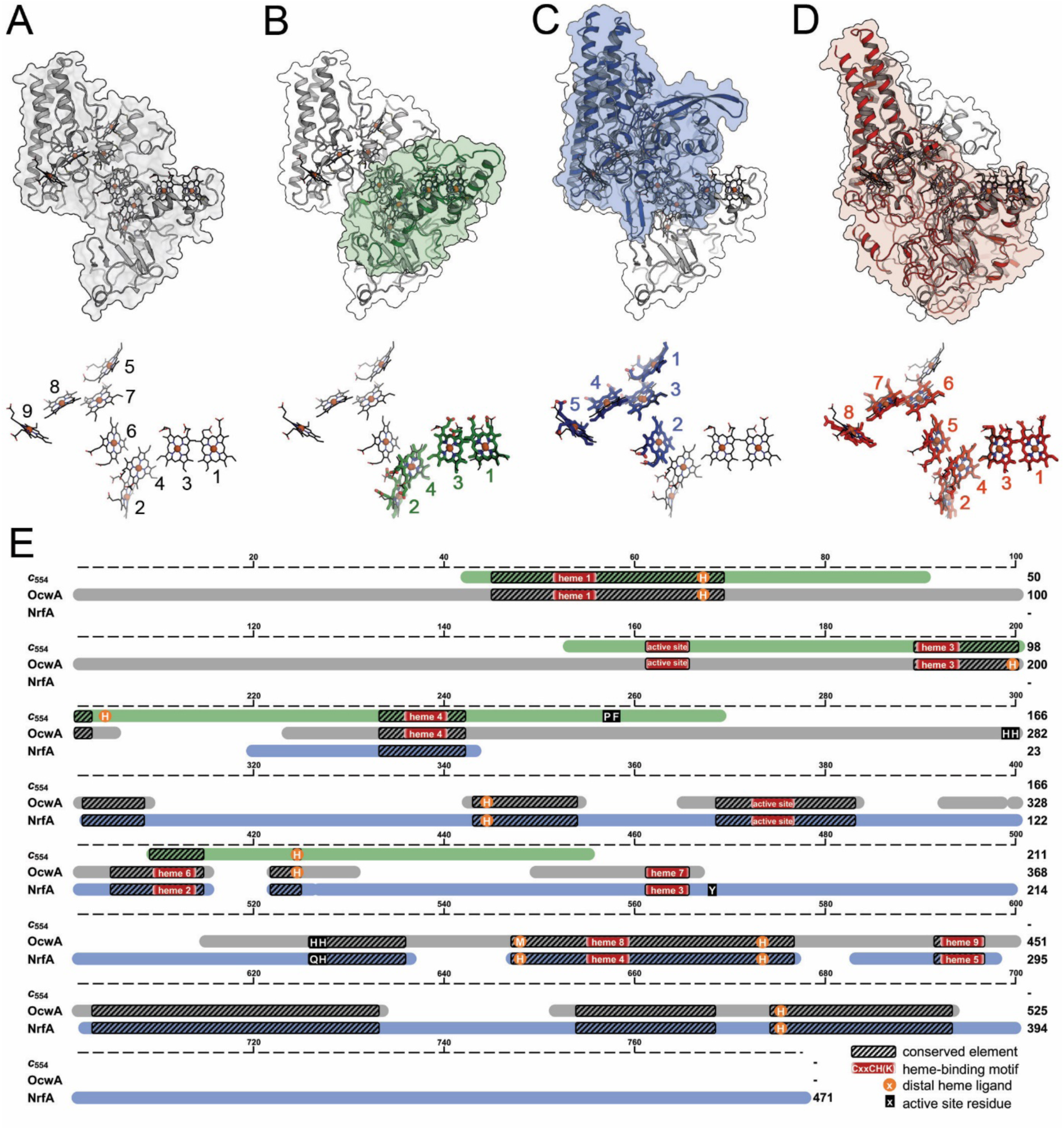
Structural and sequence similarities within the NrfA family of MHC. **A**. Structure of the OcwA monomer and heme arrangement (below), with numbering of cofactors according to their occurrence in the protein sequence. **B**. Structural alignment with *Nitrosomonas europaea* cytochrome *c*_554_ (PDB 1FT5) showing the heme numbering of this cytochrome. **C**. Structural alignment with *Wollinella succinogenes* NrfA (PDB 1FS7) showing the heme numbering of this pentaheme cytochrome. **D**. Structural alignment with *W. succinogenes* MccA (PDB 4RKM) showing the numbering of this octaheme cytochrome. **E**. Schematic presentation of a structure based amino acid sequence alignment of OcwA, NrfA and c_554_ highlighting axial heme ligands (orange circles) and distal residues of the active site(s) (black boxes). Hatched boxes represent areas with high structural homology. Heme motifs are shown with red boxes.

### Electron transfer in OcwA

The redox behavior of OcwA was explored by cyclic voltammetry. The experimental configuration used closely mimics the physiological context of this protein that is located at the surface of *T. potens* JR and exchanges electrons with electrodes in MFCs. OcwA formed a stable film at the surface of the pyrolytic graphite “edge” electrode and displayed reversible electrochemistry over a wide potential window ranging from +100 mV (fully oxidized) to −450 mV (fully reduced) *vs.* SHE (Figure 5A). This potential window is similar to that previously observed for other terminal reductases present at the cell surface of other electroactive organisms (23). This range of potentials shows that OcwA may function as the terminal reductase of *T. potens* JR, being responsible to transfer electrons to the electrode. Indeed, the overall rate of interfacial electron transfer between OcwA and the electrode determined from trumpet plots (Figure SM4) is approximately 150 s^−1^, which is in the same range as those reported for other outer-membrane terminal reductases (∼100 s^−1^) (24). Moreover, kinetic experiments with amorphous iron oxide, which is known to sustain the growth of *Thermincola*, showed that OcwA has the necessary reactivity to act as a terminal reductase in this organism (Figure SM5).

**Figure 5.**
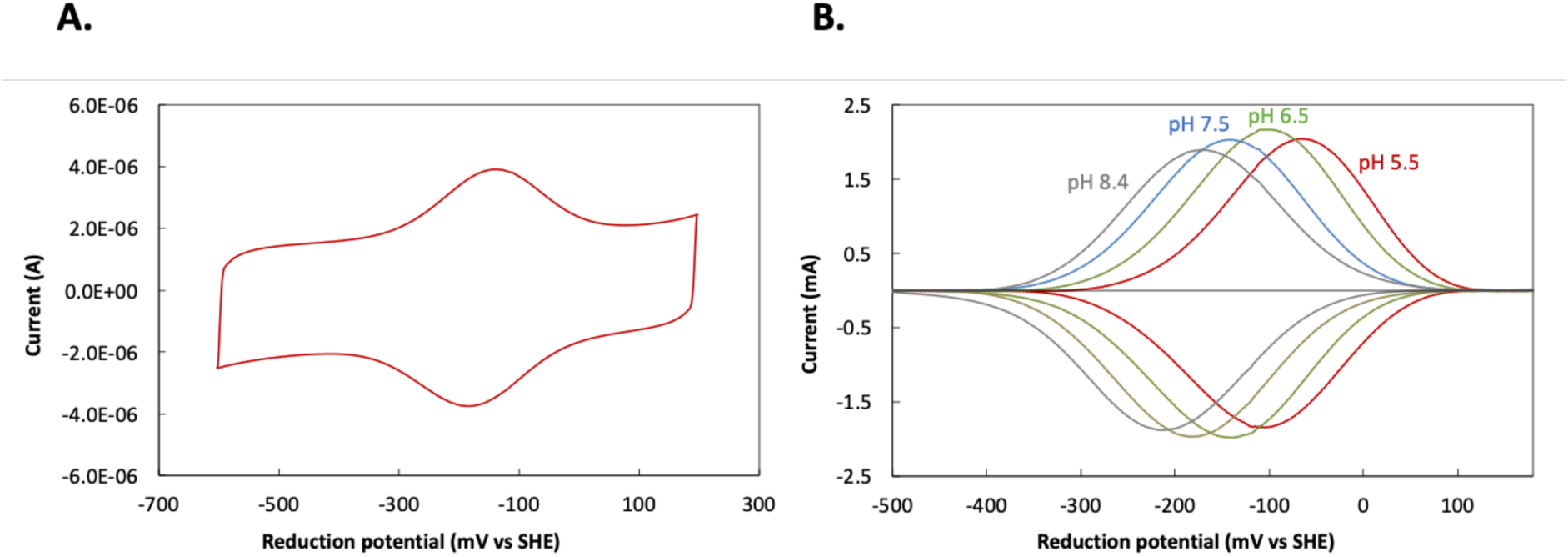
Cyclic voltammetry of OcwA. **A.** Raw voltammogram obtained at a scan rate of 200 mV/s at pH 7.5. **B.** Baseline-subtracted data of the voltammograms obtained at a scan rate of 200 mV/s at different pH values.

Electrochemical titrations of OcwA followed by cyclic voltammetry, showed pH dependence of the signals (Figure 5B), which is an indication that electron transfer is coupled to proton transfer in the physiological range (i.e. redox-Bohr effect) (25, 26). This allows redox proteins with multiple closely packed centers the possibility to become fully reduced, by balancing the electrostatic repulsion that would arise from up taking multiple negative charges with the uptake of protons.

Since *T. potens* JR was shown to use soluble electron shuttles (27), we tested the ability of OcwA to interact with anthraquinone-2,6-disulfonate (AQDS), flavin mononucleotide (FMN), riboflavin (RF) and phenazine methosulfate (PMS). These electron shuttles are typically found in environments associated with electroactive microorganisms and AQDS is known to support the growth of *T. potens* JR (10). Stopped-flow experiments showed that OcwA is oxidized by the four electron shuttles tested (Figure 6A). The degree of oxidation of OcwA at the end of the experiment is indicative of a thermodynamic equilibrium between the protein and the electron shuttles, where the different endpoints observed correlate with the midpoint reduction potential of the various electron shuttles (Figure 6B).

**Figure 6.**
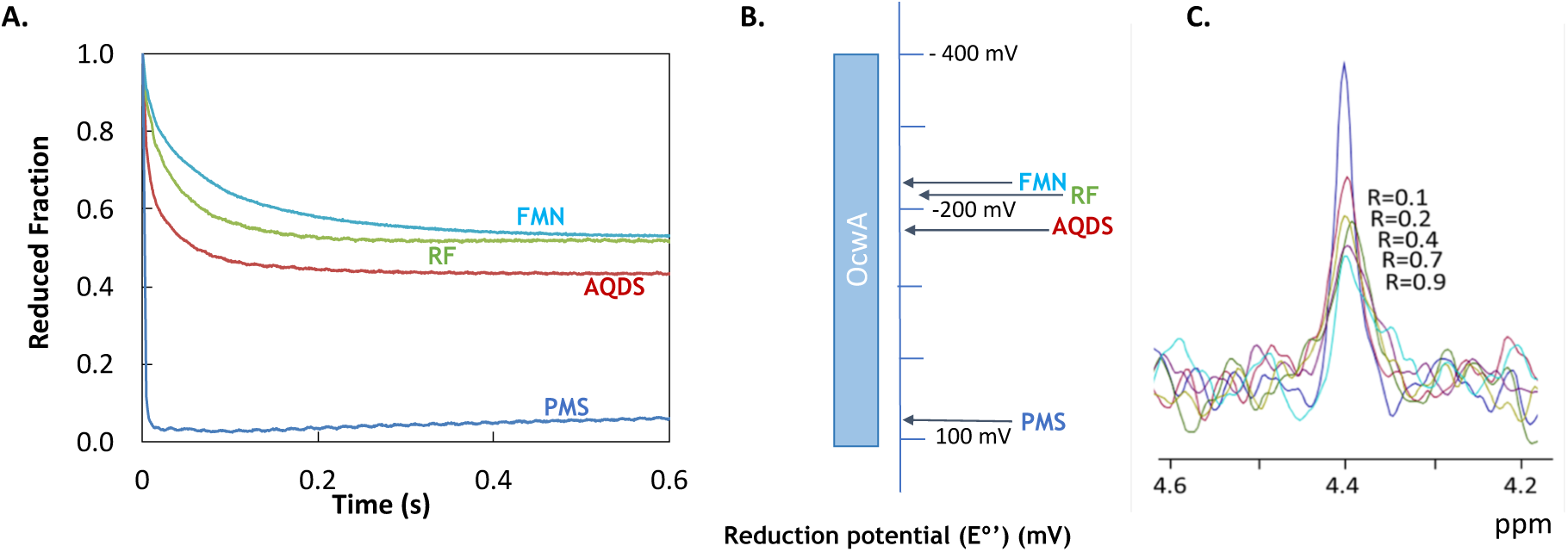
Reactivity of OcwA with electron shuttles. **A.** Kinetic data obtained by mixing OcwA (0.43 μM) with excess of AQDS (36 μM), PMS (9 μM), FMN (19 μM) and RF (14 μM). **B.** Midpoint reduction potentials of the different electron shuttles versus the redox-active range of OcwA at pH 7.6 **C.** ^31^P 1D NMR spectra of FMN in the presence of increasing amounts of OcwA obtained at 25 <u>°</u>C.

To assess the binding of the electron shuttles to OcwA we used ^1^H NMR spectroscopy. The comparison of the high-frequency region spectra obtained for OcwA alone and after addition of increasing amounts of the electron shuttles revealed no perturbation of the methyl signals located outside of the protein envelope arising from low-spin hemes (Figure SM6). This indicates that the binding of the electron shuttles may occur away from these methyl groups. This is however in contrast with the observation for the outer-membrane MHCs from *Shewanella* where the binding of soluble redox shuttles to MtrC, OmcA and UndA caused a disturbance of the heme methyl signals (28).

Taking advantage of the phosphorous atom contained in FMN, ^31^P NMR experiments were used to monitor the binding of this redox shuttle to OcwA. With increasing amounts of OcwA, we observed that the FMN phosphorous signal broadens and diminishes in intensity (Figure 6C). These spectral changes arise from binding of FMN to OcwA in slow exchange regime on the NMR timescale, which suggests a strong binding. This behavior is also different from what was previously reported for the outer-membrane MHCs from *S. oneidensis* MR-1, where the binding was transient and occurred in the fast regime on the NMR timescale (28).

### Catalytic activity of OcwA

Prompted by the structural homology with the NrfA family of proteins, the reactivity with nitrite, hydroxylamine and sulfite was tested, given that these are the most prevalent substrates of this type of enzymes (Figure SM7). It was observed that OcwA can reduce nitrite, but not sulfite in the conditions tested. Reduction and oxidation of hydroxylamine was also observed.

For nitrite values of V_max_ of 4 µmol/min.mg and K_M_ of 65 µM were obtained, which are significantly lower than those reported for the *bona fide* nitrite reductase pentaheme NrfA (29). The lack of reduction of sulfite, may be explained by the lack of a copper atom, that although not crucial for sulfite reduction activity (30), is present in the active site of the homologous MccA (20).

## Discussion

The importance of MHCs in extracellular electron transfer pathways of electroactive organisms has made their structural and functional characterization a priority in the bioelectromicrobiology field. This knowledge is even more significant for terminal reductases that are present at the cell surface of these organisms, where the knowledge on their mode of action is important to optimize the electron transfer process at the microbe-electrode interface (31, 32).

Surface exposed OcwA is proposed to be the terminal reductase in *T. potens* JR extracellular electron transfer pathway, where it bridges the contact between bacterial metabolism and electrodes in MFC (8). BLAST P sequencing revealed an OcwA homolog in *T. ferriacetica* (Tfer_3193) with which it shares 99% of identity (520/525 amino acids) including heme binding motifs and axial ligands (Figure SM8), thus suggesting a similar role between both proteins from different species of *Thermincola* sp. Given that a genetic system has not been developed for *Thermincola*, the generation of knock-out strains that would confirm this hypothesis is still not possible. Nonetheless, here, we have shown that OcwA from *T. potens* JR fulfills the role of a terminal reductase for extracellular electron transfer. It is capable of exchanging electrons with electrodes at similar rates to those of the surface exposed cytochromes of *Shewanella*, previously characterized, and to reduce iron oxides.

The redox-Bohr effect presented by OcwA appears to match the needs of *Thermincola* when growing as a biofilm at the surface of an electrode. As demonstrated by Lusk and co-workers upon studying the rate-limiting enzymatic response responsible for the electrochemical signal of *T. ferriacetica* (33), one of the key factors for the stability of electrochemically active biofilms at the surface of electrodes is the efficient dissipation of pH gradients to maintain the viability of the cells (34) and therefore the current generation (12). A similar proton-electron coupling is also found in the cell-surface exposed MHCs from *Shewanella* (23, 35). Also in similarity with the MHCs from *Shewanella*, OcwA shows an internal organization of the hemes that is neither globular nor linear but instead branched. This suggests that for the MHCs found at the surface of electroactive bacteria, having multiple hemes that can work as entry and exit electron points is an important design feature.

It is, however, at the level of the differences with previously characterized MHCs found at the surface of *Shewanella* that OcwA expands our current understanding of nature to efficiently perform extracellular electron transfer. OcwA shows for the first time in cell-surface MHCs a variety of heme coordination environments, including the open distal axial coordination position of hemes 2 and 5. High-spin hemes are typically associated with active sites where substrates are bound, and chemical reactions take place. Interestingly, the heme-binding motif of the high-spin hemes 2 and 5 in OcwA are not identical to the catalytic heme of most known NrfA proteins (i.e. CXXCK motif), nor to the P460 catalytic heme of hydroxylamine oxidising HAO, even though OcwA can perform both reactions. Although, organisms from *Thermincola* sp. were never reported to grow in the presence of nitrite or hydroxylamine (10), the ability of OcwA to reduce these substrates may be advantageous for cell survival in anaerobic environments. Indeed, OcwA may work as a detoxifying enzyme when these organisms encounter these compounds, which is a novelty for cell-surface terminal reductases. Furthermore, OcwA interacts with soluble electron shuttles, namely FMN in a distinct way of its counterparts from S. *oneidensis*, OmcA and MtrC whose interaction has been proposed to be transient and modulated by redox active disulfide bridges (33, 36). In fact, OcwA not only lacks such bridges but also binds strongly to FMN.

Finally, rather than being a truncated member of the MtrC/OmcA/UndA protein family lacking one heme, OcwA is structurally related to NrfA, cytochrome *c*_554_ and MccA, which are key proteins in the nitrogen and sulfur cycles. It was proposed that the octaheme cytochrome *c* family evolved from duplication of an ancestral *nrfA* gene in delta-proteobacteria, which was then extended by fusion with a triheme cytochrome *c* (21). The structure and catalytic versatility of OcwA provides a more nuanced view of this evolutionary storyline, now in the context of Gram-positive bacteria. From the structural point of view, OcwA can be described as a pentaheme cytochrome fused to the tetrahemic scaffold of cytochrome *c*_554_. Indeed, cytochromes of the NrfA, *c*_554_ and octaheme families are evolutionarily related (2, 21). The structure of OcwA raises the possibility that the extant octaheme cytochromes evolved by the loss of heme 2 (HAO) or heme 5 (MccA) from ancestral nonaheme cytochromes similar to OcwA.

In conclusion, the structural and functional characterization of OcwA reveals that this terminal reductase for extracellular electron transfer is unrelated to the known terminal reductases, MtrC, OmcA and UndA. This indicates that the process of extracellular electron transfer evolved independently more than once to accommodate the catabolic needs of microorganisms living in environments with access to solid terminal electron acceptors. Moreover, the particular case of OcwA reveals a multifunctional link between the biogeochemical cycles of nitrogen and iron. Indeed, being able to react with electrodes, N-oxides, iron and small molecule redox shuttles, OcwA may work as the respiratory ‘swiss-army knife’ of *Thermincola* allowing this Gram-positive bacterium to grow and survive in environments with variable access to solid and soluble electron acceptors.

## Materials and Methods

### Protein production and purification

The OcwA protein was produced according to the literature (37) with minor changes using the primers listed in Table 1. Briefly, a chimeric gene, containing the signal peptide of small tetraheme cytochrome c (*stc*) from *S. oneidensis* MR-1 fused with the gene sequence of *therjr_2595* without native signal peptide was constructed. *E. coli* JM109 (DE3) was co-transformed with this chimeric gene previously cloned into pBAD202/D-TOPO vector (38) and plasmid pEC86 that contains the cytochrome *c* maturation system (39). Transformed *E.coli* strain were cultured in TB medium with 50 µg kanamycin and 35 µg of chloramphenicol at 37°C for 6 hours. At mid log phase (∼6 h after inoculation), the temperature of the growing culture in TB medium previously set at 37°C was lowered to 30^°^C. The cells were allowed to grow for additional 18 h and were then pelleted by centrifugation at 10,000×*g* for 10 min at 4°C and resuspended in osmotic shock solution (0.2 M Tris-HCl (pH 7.6) with 0.5 M sucrose, 0.5 mM EDTA and 100 mg/L lysozyme) with protease inhibitor (Sigma) and DNase (Sigma) using a ratio of 200 ml of osmotic shock solution for each 8L of culture media. The spheroplast solution was incubated at 4°C for 40 min with gentle stirring. The periplasmic fraction containing the recombinant protein was cleared by ultracentrifugation at 200,000×*g* for 1h at 4°C and dialyzed overnight against 20 mM Tris-HCl (pH 7.6). The dialyzed protein extract was purified *via* a His-trap column (GE Healthcare) previously equilibrated with 20 mM sodium phosphate buffer, 20 mM imidazole and 500 mM NaCl (pH 7.6). To elute the bound protein, a single step was used with 20 mM sodium phosphate buffer, 300 mM Imidazole and 500 mM NaCl (pH 7.6). The resulting fraction was dialyzed overnight against 20 mM Tris-HCl (pH 7.6). A diethylaminoethyl (DEAE)-biogel column (Biorad) pre-equilibrated with 20 mM Tris-HCl (pH 7.6) was used as a final purification step. OcwA was eluted in 20 mM Tris-HCl buffer with 120 mM NaCl (pH 7.6). The purified protein was dialyzed against 20 mM potassium phosphate buffer (pH 7.6) with 100 mM KCl. SDS-PAGE (12% resolving gel) and UV-visible spectroscopy were used after each purification step to select the fractions containing the target protein and to evaluate its purity. The purified protein had an absorbance ratio (A_Soret peak_/A_280nm_) larger than 5.0.

**Table 1.**
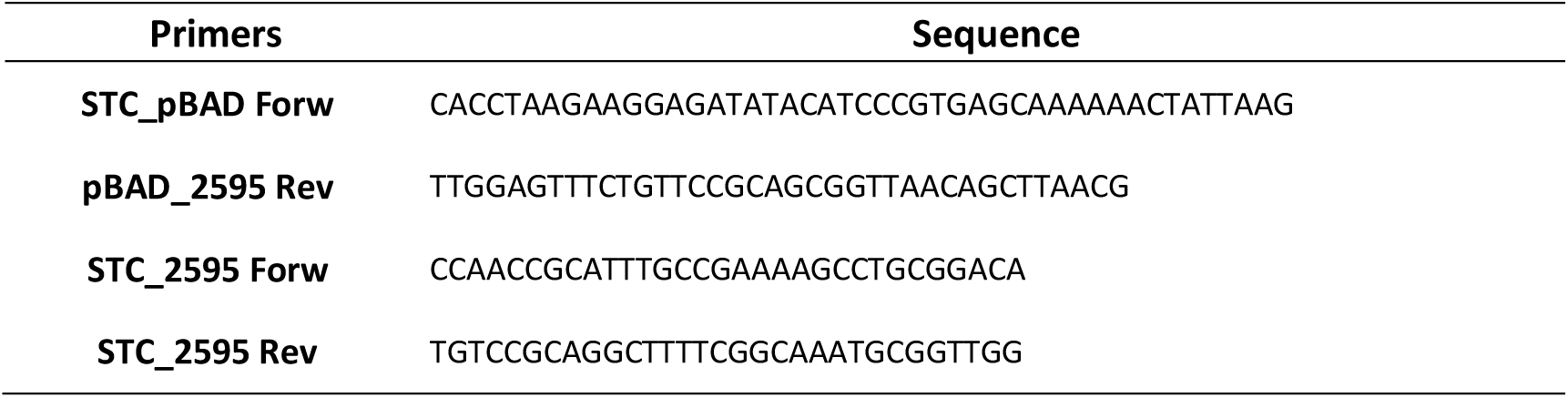
Oligonucleotides used to construct the chimeric gene

### Crystallization and structure determination

For structure determination, crystals were grown in an anoxic chamber (Coy) under N_2_/H_2_ atmosphere (95/5 vol %) at room temperature. Sitting drop vapor diffusion experiments were set up by adding 0.5 µL of protein solution (6 mg/mL) to 0.5 µL of reservoir solution. Well-diffracting crystals were obtained in conditions containing 24-27% polyethylene glycol 3350, 0.1 M HEPES, pH 7-7.5 and 0.1-0.2 M MgCl_2_ only after seeding. The crystals were harvested using 10% 2,3-butanediol as cryoprotectant in mother liquor and flash frozen in liquid nitrogen. Datasets were collected at the Swiss Light Source, Paul-Scherrer-Institut (Villigen, Switzerland) on beamline X06DA with a Pilatus 2M Detector (Dectris). Initial phase information was obtained by single anomalous diffraction with a dataset collected at the iron edge, *λ* = 1.729 Å, using the PHENIX suite (40) for automatic phasing and initial model building. An initial structural model was refined in iterative steps using REFMAC5 (41) and COOT (42) to 2.7 Å resolution. This solution was then used for molecular replacement with MOLREP (43) as part of the CCP4 suite (44) for the more highly resolved structure at 2.2 Å. The final model comprised residues 42 to the very C-terminus at residue 525 in chain A, and residues 43 to 525 in chain B. Coordinates and structure factors have been deposited with the protein data bank under the accession code 6I5B. Data collection and refinement statistics are summarized in supplementary table SM1.

### NMR experiments

For all the NMR experiments, OcwA prepared in 20 mM potassium phosphate buffer (pH 7.6) with 100 mM KCl was lyophilized and resuspended in D_2_O. An excess of sodium dithionite was used to reduce the protein. ^1^H NMR experiments were performed on a Bruker Avance II 500 MHz spectrometer equipped with a QXI probe for ^1^H detection, and a SEX probe for ^31^P detection. All NMR data were processed in the Topspin 3.2 software. ^1^H NMR spectra were acquired before and after lyophilization to ensure that protein integrity was preserved.

To study the influence of the electron shuttles on OcwA, NMR experiments were performed as previously described (28) using antraquinone-2,6-disulfonate (AQDS), flavin mononucleotide (FMN), riboflavin (RF) and phenazine methosulfate (PMS). Stock solutions of the different electron shuttles were prepared in 20 mM potassium phosphate buffer (pH 7.6) with 100 mM KCl. ^1^H NMR spectra performed at 25°C on a Bruker Avance II 500 MHz NMR spectrometer equipped with TCI cryoprobe for ^1^H detection were acquired before and after the addition of the electron shuttles (molar ratios 0.5:1, 1:1 and 3:1 electron shuttle:protein). For the ^31^P-NMR binding experiments, samples containing 100 µM of FMN prepared in 20 mM phosphate buffer (pH 7.6) with 100 mM KCl were titrated against increasing concentrations of OcwA at 25°C.

### EPR experiments

The OcwA solution was prepared in 20 mM potassium phosphate buffer (pH 7.6) with 100 mM KCl to a final concentration of 200 μM. The reduced state was obtained by addition of an excess of sodium borohydride. EPR spectra were recorded on a Bruker ESP 380 spectrometer equipped with an ESR 900 continuous-flow helium cryostat (Oxford Instruments). Temperature: 7 K; microwave frequency: 9.39 GHz; modulation amplitude: 1.0 mT; microwave power: 2 mW.

### Cyclic Voltammetry of OcwA

The electrochemical setup was assembled as previously described (45, 46). Electrochemical measurements were performed in an anaerobic glovebox (JACOMEX, France) with a nitrogen atmosphere (residual O_2_ < 1 ppm). A pyrolytic graphite edge electrode (PGE surface approx. 3 mm^2^) was polished with alumina slurry (Buehler, 1μm) and then coated with 0.5 μl of a solution of OcwA (stock solution previously diluted in 20 mM potassium phosphate buffer (pH 7.6) with 100 mM KCl to a final concentration of 50 μM) and left to dry for approximately 5 min. The buffers used in the electrochemical experiments were prepared by mixing 5 mM of HEPES, MES and TAPS and 100 mM KCl. The desired pH values were adjusted with 1 M NaOH or HCl solutions. Experiments were performed at 25°C using an electrochemical cell consisting of an Ag/AgCl (saturated KCl) reference electrode in a Luggin sidearm and a platinum wire counter electrode. Cyclic voltammetry was performed with an Autolab electrochemical analyzer (PGSTAT-128N) with an analogue scan generator controlled by GPES software. The potentials are quoted with reference to standard hydrogen electrode (SHE) by addition of 0.197 V to those measured (47). The electrochemical data were analyzed using the QSOAS program available at www.qsoas.org (48).

### Kinetic experiments between OcwA and electron shuttles

Protein oxidation experiments were performed according to Paquete *et al.* 2014 (28) in a stopped-flow apparatus (SHU-61VX2 from TgK Scientific) placed inside an anaerobic chamber (M-Braun 150) containing less than 5 ppm of oxygen. The concentration of protein, approximately 0.4 µM after mixing was determined for each experiment by UV-visible spectroscopy using ε_409nm_ of 125.000 M^−1^ cm^−1^ per heme for the oxidized state of the protein (46, 49). Stock solutions (5 mM) of the electron shuttles antraquinone-2,6-disulfonate (AQDS), flavin mononucleotide (FMN), riboflavin (RF) and phenazine methosulfate (PMS) were prepared by dissolving weighted amounts of solid reagents in 20 mM potassium phosphate buffer (pH 7.6) with 100 mM KCl. Dilutions of the electron shuttles were prepared in degassed buffer, and their concentrations were determined by UV-visible spectroscopy using the following extinction coefficients ε_326nm_ = 5200 M^-1^ cm^-1^ for AQDS (50), ε_445nm_ = 12,200 M^-1^ cm^-1^ for FMN (51), ε_445nm_ = 12,500 M^-1^ cm^-1^ for RF (52), and ε_387nm_ = 26,300 M^-1^cm^-1^ for PMS (53). Reduced OcwA was obtained by mixing the protein with small volumes of concentrated sodium dithionite solution. UV-visible spectroscopy was used to confirm that there was no excess of dithionite using ε_314nm_ =8.000 M^−1^ cm^−1^ (54). Oxidation by electron shuttles was monitored by measuring the absorption changes at 552 nm upon mixing reduced OcwA with each of them. The temperature of the kinetic experiments was maintained at 25°C using an external circulating bath (28).

### Catalytic experiments with OcwA

The enzymatic reduction of sulfite, nitrite and hydroxylamine by OcwA was tested at 55°C in an anaerobic chamber (Coy). All the assays were performed in 50 mM potassium phosphate (KPi) buffer at pH 7.0, with 1 mM methyl viologen previously reduced with zinc granules as described in the literature (30). The reaction was started by the addition of the substrate, followed by OcwA, and the reaction was monitored by the oxidation of methyl viologen at 732 nm. Sodium sulfite concentrations were tested in the range of 100 µM to 6 mM, while sodium nitrite was tested in the range 20 µM to 150 µM, and hydroxylamine in the range 10 µM to 200 µM. The final concentrations of OcwA ranged from 50 to 400 nM measured prior to the experiments using ε_409_ nm of 125.000 M^−1^ cm^−1^ per heme for the oxidized state of the protein.

The enzymatic oxidation of hydroxylamine by OcwA was tested using 20 µM of phenazine methosulfate (PMS) and 400 µM of 3-(4,5-dimethyl-2-thiazolyl)-2,5-diphenyl-2H-tetrazolium bromide (MTT) as electron acceptors (19). The reaction was started by the addition of hydroxylamine (75 µM to 500 µM), followed by 100 µM OcwA, and the reduction of MTT was monitored at 578 nm.

The enzymatic reduction of amorphous iron (III) oxide by OcwA was also performed at 55°C in the same anaerobic chamber. Amorphous iron (III) oxide was prepared as described in (11), a solution of FeCl_3_ was titrated with 10% (w/v) NaOH (pH 11) and brown precipitate was formed. In each assay iron (III) oxide was added to 488 nM OcwA, previously reduced with sodium dithionite, in 50 mM potassium phosphate buffer (pH 7.0). Iron (III) oxides were tested in the range of 14.3 to 55.5 mg/L. UV-visible spectra of OcwA were obtained as isolated (oxidized state), after the addition of sodium dithionite (reduced state), and after addition of the iron oxides. All solutions were previously degassed.

## Acknowledgements

The authors are grateful to Dr Christophe Léger for helpful comments and suggestions. This work was supported by Fundação para a Ciência e a Tecnologia (FCT) Portugal [PTDC/BBB-BQB/4178/2014, and PTDC/BIA-BQM/30176/2017], by Project LISBOA-01-0145-FEDER-007660 (Microbiologia Molecular, Estrutural e Celular) funded by FEDER funds through COMPETE2020 - Programa Operacional Competitividade e Internacionalização (POCI), and by ITQB research unit GREEN-it “Bioresources for sustainability” (UID/Multi/04551/2013). This work has received funding from the European Union’s Horizon 2020 research and innovation programme under grant agreement No 810856. The NMR spectrometers at CERMAX are part of the National NMR Network (PTNMR) and are partially supported by Infrastructure Project No 022161 (co-financed by FEDER through COMPETE 2020, POCI and PORL and FCT through PIDDAC). N-terminal data was obtained by the N-terminal Sequencing Facility at ITQB.

